# Segmentation aware probabilistic phenotyping of single-cell spatial protein expression data

**DOI:** 10.1101/2024.02.29.582827

**Authors:** Yuju Lee, Edward L. Y. Chen, Darren C. H. Chan, Anuroopa Dinesh, Somaieh Afiuni-Zadeh, Conor Klamann, Alina Selega, Miralem Mrkonjic, Hartland W. Jackson, Kieran R. Campbell

## Abstract

Spatial protein expression technologies can map cellular content and organization by simultaneously quantifying the expression of >40 proteins at subcellular resolution within intact tissue sections and cell lines. However, necessary image segmentation to single cells is challenging and error prone, easily confounding the interpretation of cellular phenotypes and cell clusters. To address these limitations, we present STARLING, a novel probabilistic machine learning model designed to quantify cell populations from spatial protein expression data while accounting for segmentation errors. To evaluate performance we developed a comprehensive benchmarking workflow by generating highly multiplexed imaging data of cell line pellet standards with controlled cell content and marker expression and additionally established a novel score to quantify the biological plausibility of discovered cellular phenotypes on patient derived tissue sections. Moreover, we generate spatial expression data of the human tonsil – a densely packed tissue prone to segmentation errors – and demonstrate cellular states captured by STARLING identify known cell types not visible with other methods and enable quantification of intra- and inter- individual heterogeneity. STARLING is available at https://github.com/camlab-bioml/starling.

## Introduction

Highly multiplexed imaging technologies such as Imaging Mass Cytometry (IMC)^1^ are a set of state-of-the-art assays that allow simultaneous quantification of up to approximately 40 protein targets at sub-cellular resolution in two dimensions^2^. This enables delineation of high dimensional single-cell phenotypes and cellular communities in both archival patient tissue sections as well as tissues from animal models and cell lines. Together, their applications have uncovered cellular phenotypes and spatial architectures that stratify patient outcomes and predict disease course across a range of pathologies including breast^3,4^, lung^5^, melanoma^6^, and brain^7^ cancers, along with other pathologies such as diabetes^8^.

A major step in the analysis of multiplexed image cytometry is the process of cell segmentation that identifies where in the original (pixel-level) data cells are. Commonly, the data is subsequently summarized to the single-cell level by taking the mean expression of markers over all pixels belonging to a given cell along with cell size and/or morphology information and applying standard data analysis pipelines developed for suspension cytometry such as visualization^9^, clustering^10,11^, and cell type assignment^12^. For the process of segmentation, multiple algorithms have been proposed ranging from hand-crafted machine learning pipelines^3^ to deep learning methods including CellPose^13^ and Mesmer^14^.

However, it is increasingly being recognized that cell segmentation is an inherently error-prone process due to resolution limitations, three-dimensional tissue architecture with cells overlapping in the z-plane, and algorithmic imperfections. These issues result in signal spillover between adjacent cells^15,16^, over-segmentation resulting in doublets, and cellular projections being mis-assigned^17^. Upon summarization to the single-cell level, this results in cellular phenotypes that are frequently the average of two or more actual cells, which in turn leads to cells or clusters that co-express implausible protein expression combinations. For example, CD3 and CD20 should be exclusively expressed on T and B lymphocytes respectively^18^, but cell populations that co-express them are frequently found in imaging mass cytometry datasets^3,4^.

Despite these challenges, few computational methods for highly multiplexed imaging address these limitations. Classical clustering methods for suspension CyTOF frequently re-used for multiplexed imaging^3,4^ such as PhenoGraph^11^ and FlowSOM^10^ all assume pure single cell expression as a starting point. Multiple strategies have attempted to augment resolution to improve segmentation, but segmentation errors occur even in high resolution imaging methods^19,20^. CellSighter^17^ is a recently developed method for multiplexed imaging that attempts to circumvent segmentation errors by operating on the raw imaging data with data augmentation, but is used for supervised classification of cell types given a labeled training set rather than *de novo* discovery of cell phenotypes as is standard in a clustering workflow.

Conversely, doublet detection and removal have received significant attention in single-cell transcriptomics, with multiple methods^21–24^ and benchmarking works available^25^. To-date, no methods have been proposed to learn denoised cellular phenotypes while accounting for cell segmentation errors in highly multiplexed imaging.

To help tackle this, we present STARLING (SegmenTation AwaRe cLusterING), a novel probabilistic clustering method specifically designed to account for the fact that single-cell expression profiles from multiplexed imaging may represent the combination of two or more actual cell populations. We introduce the concept of a plausibility score that quantifies the quality of a given clustering to infer biologically meaningful cellular phenotypes and introduce a further benchmarking framework to quantify the ability of methods to detect segmentation errors from multiplexed imaging. We further generate IMC data of cell pellets of controlled cell lines to quantify the ability of different methods to re-infer cellular clusters when the input cell types are both controlled and known. Finally, we generate IMC data from 16 regions of interest (ROIs) from the human tonsil across 3 donors — a secondary lymphoid organ densely packed with immune cells — and demonstrate how the STARLING immunophenotypes detect spatial single-cell communities and highlight intra- and inter-donor variability. An open source Python package implementing the STARLING method is available at https://github.com/camlab-bioml/starling.

## Results

### STARLING: a probabilistic model for segmentation error aware clustering of highly multiplexed imaging data

Multiple artifacts give rise to segmentation errors in multiplexed imaging data, including doublets where a segment covers two or more cells, spatial spillover, 3D tissue effects, and cellular projections^16,17^. The consequence is that when the error involves cells from different cell types — previously described as heterotypic doublets^24^ — clusters are formed due to the composition of different cells within a segment that are not “real” cell types in the source biological material (**Fig. 1A**). This can be shown by clustering data from multiple recent publications giving rise to cell types that co-express biologically implausible marker combinations (**S. Fig. 1**). The consequence is that outside the few scenarios where implausible clusters may be manually identified, it is often difficult to trust the cellular phenotypes identified from such experiments without follow-up validation, and even more so trust every cell label within the thousands or millions of cells in a dataset. At worst, novel cell types or spatial features may be identified that are in fact purely due to segmentation artifacts.

**Figure 1.**
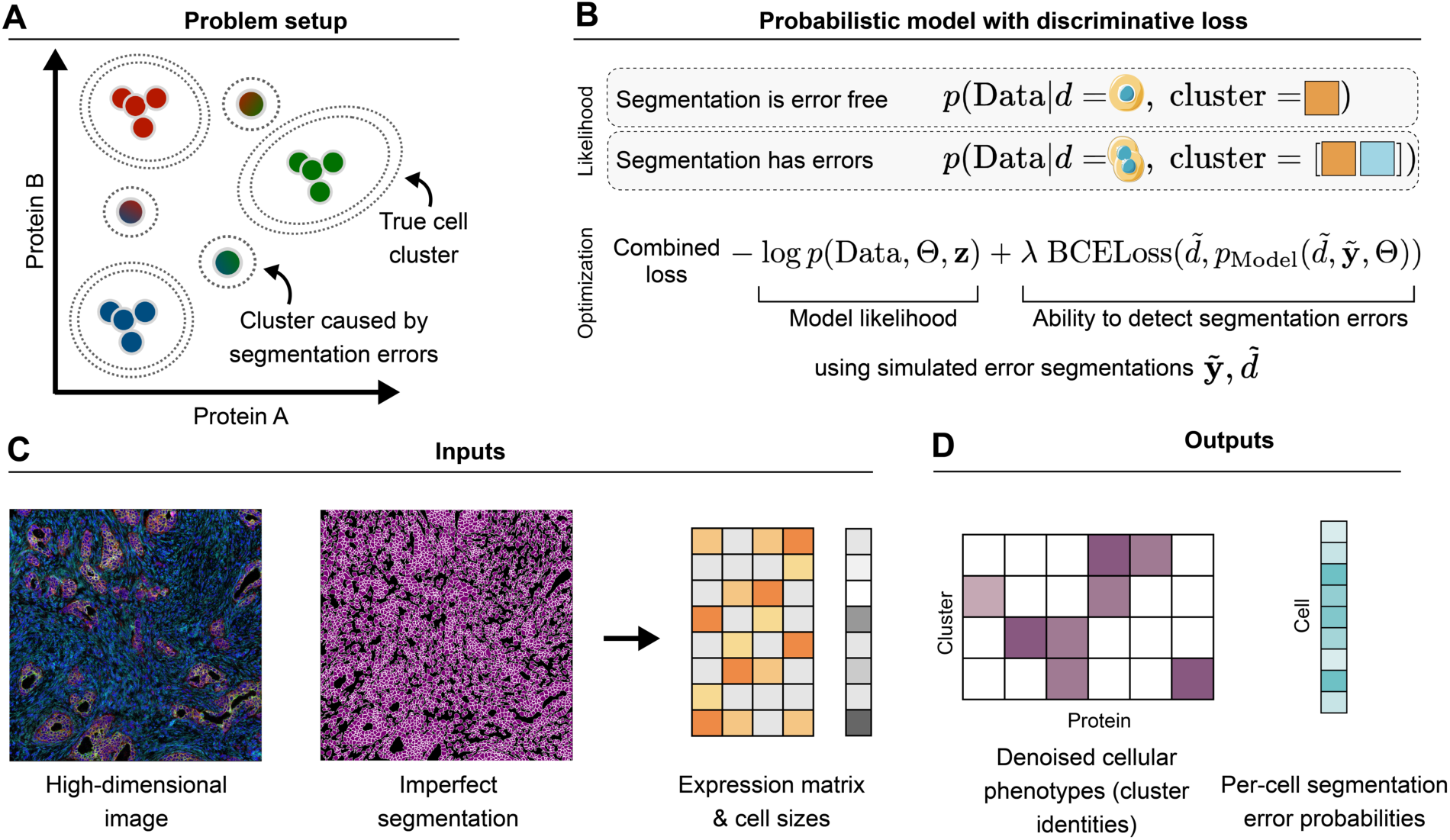
**A** Segmentation errors in highly multiplexed imaging experiments induce clusters formed by the composition of multiple true cell types. **B** Our probabilistic model STARLING for clustering highly multiplexed imaging data models whether each cell is observed segmentation error free, and if not models the composition of cell types leading to the observed cell. The loss function jointly maximies the likelihood of the data and the ability to discriminate on-the-fly simulated segmentation errors. **C** The input to STARLING includes a high dimensional multiplexed image and imperfect segmentation that is summarized to an expression matrix with cell sizes. **D** The output includes both the denoised cell types along with a per-cell estimate of segmentation errors.

To help address this, we developed STARLING (**S**egmen**T**ation **A**waRe c**L**uster**ING**), a novel probabilistic clustering method to identify true cell phenotypes in the presence of segmentation errors. STARLING specifically models a per-cell probability of a segmentation error, along with the true cluster identities of the underlying cells segmented together as part of the error. The joint likelihood of the data, cluster identities and assignments, and segmentation error probabilities are then optimized along with a discriminative loss that regularizes the model to be able to both accurately predict errors and orient which clusters are likely true compared to error-induced (**Fig. 1B** and **Methods**). As input, STARLING requires only the high-dimensional image along with imperfect cell segmentation that are summarized to a per-cell expression profile along with physical cell size in pixels (**Fig. 1C**) as is commonly output by a range of pipelines for highly multiplexed imaging data^26,27^. After optimization of the loss, STARLING outputs both the denoised cluster identifies representing the cellular phenotypes of the underlying cells in the input biological material, as well as a per-cell segmentation error probability that may be useful for follow-up visualization and analysis (**Fig. 1D**).

### Scoring the biological plausibility of clusters inferred from multiplexed images

We first sought to validate that the clusters returned by STARLING showed more biologically realistic cellular phenotypes than existing strategies. Given true cell types are *a priori* unknown and benchmarking using simulated data can introduce unrealistic assumptions, we built upon previous work^16,28^ that uses the co-expression of marker proteins to evaluate the quality of the inferred cell clusters. Specifically, we generalized this idea to create the *plausibility score* (**Fig. 2A**) that takes an *a priori* known set of protein pairs that either exhibit mutually exclusive expression or conditional co-expression. For example, the T lymphocyte marker CD3 should never be co-expressed with the B lymphocyte marker CD20; conversely, CD3 should never be expressed in the absence of the pan-leukocyte marker CD45. By curating a list like this for each dataset’s antibody/antigen panel, for a given clustering we can simply count the proportion of centroids that fall in the expected regions using a given threshold, and designate a normalized version of this as the plausibility score (**Methods**).

**Figure 2.**
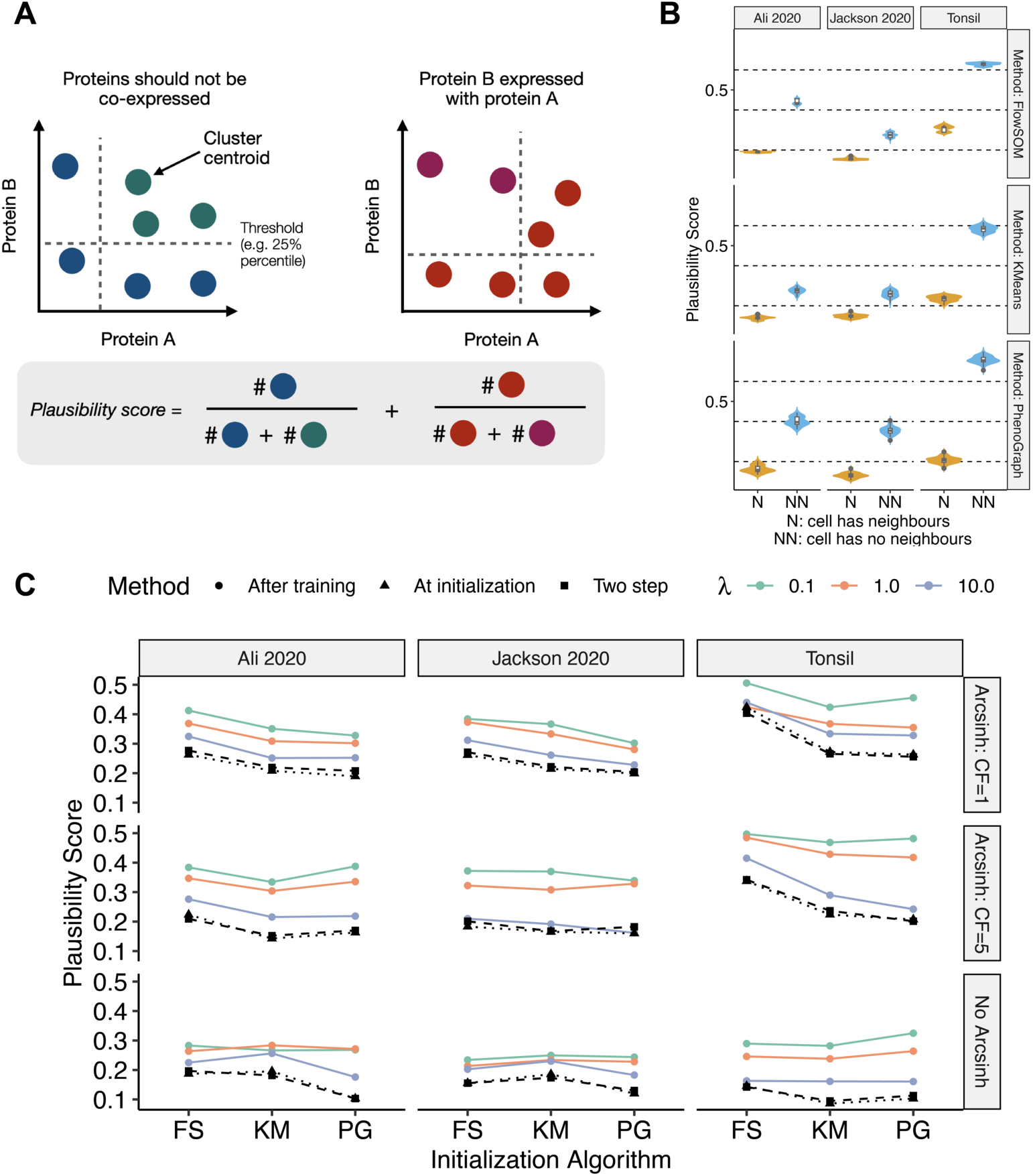
**A** Pairs of proteins known to not be co-expressed (left) or show conditional expression (right) are collated for each dataset, and the *plausibility score* defined as the proportion of centroids that lie in the expected regions under some threshold of expression. **B** To validate the plausibility score on real data, we took three IMC datasets and separated cells into those that have neighbours (N) or no neighbours (NN). We clustered each with three clustering algorithms and computed the plausibility score on the centroids. Cells with no neighbours and therefore likely fewer segmentation errors show significantly higher plausiblity scores than those with neighbours. **C** The plausibility score quantified across three datasets, three normalization scenarios, and three hyperparameter settings. The black triangles represent the score at initialization using three popular clustering algorithms (FS: FlowSOM, KM: K-means; PG: PhenoGraph) and coloured lines after training. The black squares represent the two step approach of first removing likely doublets.

To validate this plausibility score we exploited the fact that cells isolated with no neighbours should have fewer segmentation errors than those in densely packed regions with many neighbours. Using three common clustering algorithms (FlowSOM^10^, PhenoGraph^11^, Kmeans) we clustered three multiplexed imaging datasets — two published breast cancer atlases^3,4^ and a human tonsil dataset generated for this study (see **Methods**) — separately for cells that have neighbours and those that do not. We then computed the plausibility score for each set of clustering using marker pairs derived for each dataset’s antibody panel (**S. Table 1**). Across all datasets and clustering algorithms, the plausibility score is significantly higher for cells with no neighbours than those with neighbours (**Fig. 2B**), validating its utility as an independent measure of biologically realistic cellular clusters from highly multiplexed imaging data.

Next, we benchmarked the plausibility score of clusters found by STARLING compared to the three established clustering algorithms on the three highly multiplexed imaging datasets. To comprehensively disentangle the factors associated with benchmarking performance, we initialized STARLING using each of the three clustering algorithms, and compared the initial plausibility score (equivalent to clustering using the existing methods) to the post-STARLING clustering plausibility score. In addition, we varied the λ hyperparameter in STARLING that controls the trade-off between optimizing the model likelihood and ability to detect synthetic segmentation errors as well as different data pre-processing strategies (via archsinh normalization) commonly used in IMC data analysis^12^. We also considered a two-step approach where first a random forest model is trained to detect doublets and cells with a high probability of being a doublet are removed before the data is clustered using the baseline methods (**Methods**). The results in **Fig. 2C** demonstrates that for all clustering algorithms, hyperparameter combinations, and data normalization strategies, STARLING outperforms both the baseline initializations as well as the two-step doublet removal strategy.

### Assessing the ability of STARLING to detect cell segmentation errors

Next, to assess whether segmentation errors quantified by STARLING are reliable we adopted a multi-faceted strategy. First, we exploited the fact that MESMER^14^ — a state of the art segmentation algorithm for multiplexed imaging — can perform both whole-cell and nuclear segmentation. We segmented each image in each dataset twice and designated cells where more than one nuclei was found in the nuclear-only segmentation despite only a single cell being found in the whole-cell segmentation as likely to contain a segmentation error. We then quantified STARLING’s segmentation error probability for cells likely to contain a segmentation error compared to otherwise. The results in **Fig. 3A** shows a clear increase in segmentation error probability for cells whose nuclear segmentation conflicts with the whole-cell segmentation. Note that given the multiple ways a segmentation error can occur beyond doublets we do not expect the segmentation error probability to be zero in cells where the nuclear segmentation matches the whole-cell segmentation.

**Figure 3.**
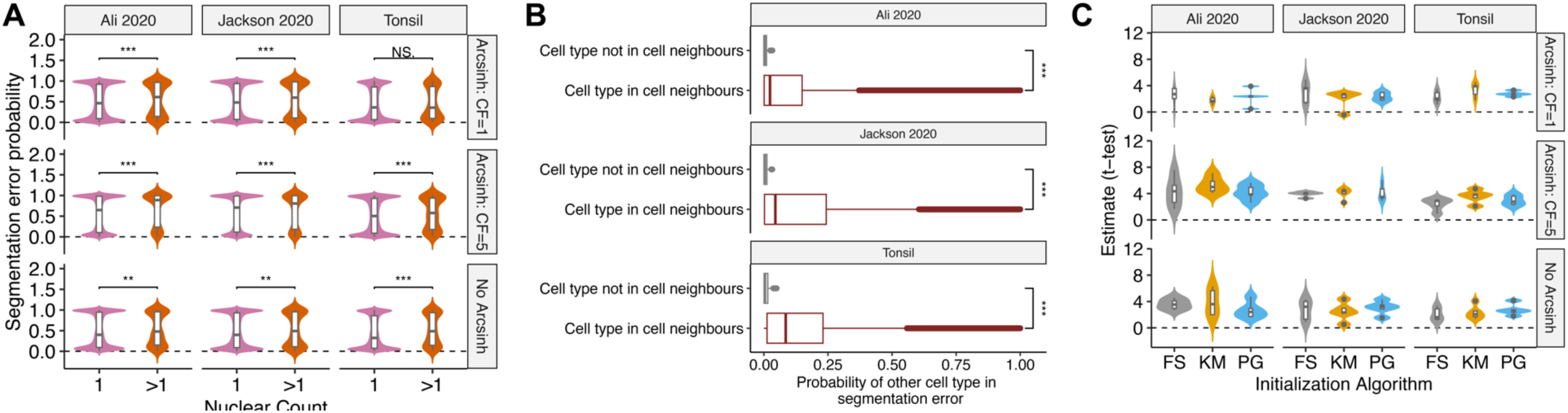
**A.** The segmentation error probability computed by STARLING based on whether each whole-cell segmented by MESMER contains one or more segmented nuclei across three datasets and three normalization schemes. **B.** The probability that the mix of cell types identified by STARLING to be involved in a segmentation error is significantly higher when those cell types involve the cell neighbours. **C.** AT-statistic comparing the convex area to total area of each cell for cells predicted to contain segmentation errors by STARLING vs those predicted to be segmentation error free.

Secondly, we quantified the ability of STARLING to detect spatial spillover despite not explicitly being given spatial information about the cells beyond size. Specifically, for each cell across the three datasets we identified the most likely cluster in the absence of segmentation errors as well as the most likely cluster of each neighbouring cell. Next, we took STARLING’s per-cell Γ matrix that quantifies the probabilities of all possible pairings of clusters that contributed to the segmentation errors involved in a given cell (**Methods**). For each cell, we quantified the value of this matrix for the pair of clusters corresponding to that cell and its neighbours and contrasted this with the pair of clusters corresponding to that cell and all clusters not present in the cell’s neighbours. We found significantly higher probability of a cluster contributing to a cell’s segmentation error when a neighbouring cell was of that cluster type than otherwise (**Fig. 3B**). This implies STARLING implicitly detects spatial spillover artifacts despite not being explicitly provided information about neighbouring cells or their expression profiles.

Finally, we reasoned that cells likely to involve segmentation errors would have altered morphology. Cell segments corresponding to doublets (i.e. two cells segmented into one) or with cellular projections will be more likely to have a concave morphology where the ratio of convex area to total area is increased compared to cells with high quality segmentations (**S. Fig. 2**). Therefore, across the three datasets, normalization strategies, and initialization algorithms, we separated cells predicted to involve segmentation errors from those predicted to be error-free and computed the per-cell ratio of convex to total cell area for each. Despite STARLING not having access to information on cellular morphology, contrasting these two values using a t-test demonstrates a significantly increased convex to total area ratio in cells predicted to have segmentation errors (**Fig. 3C**). Overall these results imply that in addition to the cellular phenotypes quantified by STARLING being biologically plausible, the segmentation error probabilities represent a robust measure of segmentation quality that may serve for useful downstream analysis and interpretation.

### Quantifying cell phenotype inference using controlled cell types

Next, we evaluated the inference of cellular phenotypes in a controlled setting. We created a cell pellet from four cell lines, each known to be positive for only one major lineage marker (CD3+ Jurkat, CD20+ RAMOS, CD68+ THP1, PanCK+ BxPC3, see **Methods**). This dataset acts as a control as each resulting cluster should be positive for only one of the major lineage markers, with any crossover due to segmentation errors. The cell lines were mixed in equal proportions, stained with an antibody panel able to discriminate different cell types (**Fig. 4A**), and subjected to IMC (**Fig. 4B**) resulting in high-dimensional quantification of the resulting cell lines with spatial resolution (**Fig. 4C**). After applying a standard analysis pipeline of segmentation using Mesmer^14^ and clustering with PhenoGraph^11^, we identified 13 distinct cell types showing heterogeneous expression profiles (**Fig. 4D**). Using a procedure to assign clusters to cell types based on single marker positivity (**Methods**), we were able to confidently assign 9/13 clusters to the cell lines of origin, with 4/13 labeled as “Unsure” due to multi-marker positivity.

**Figure 4.**
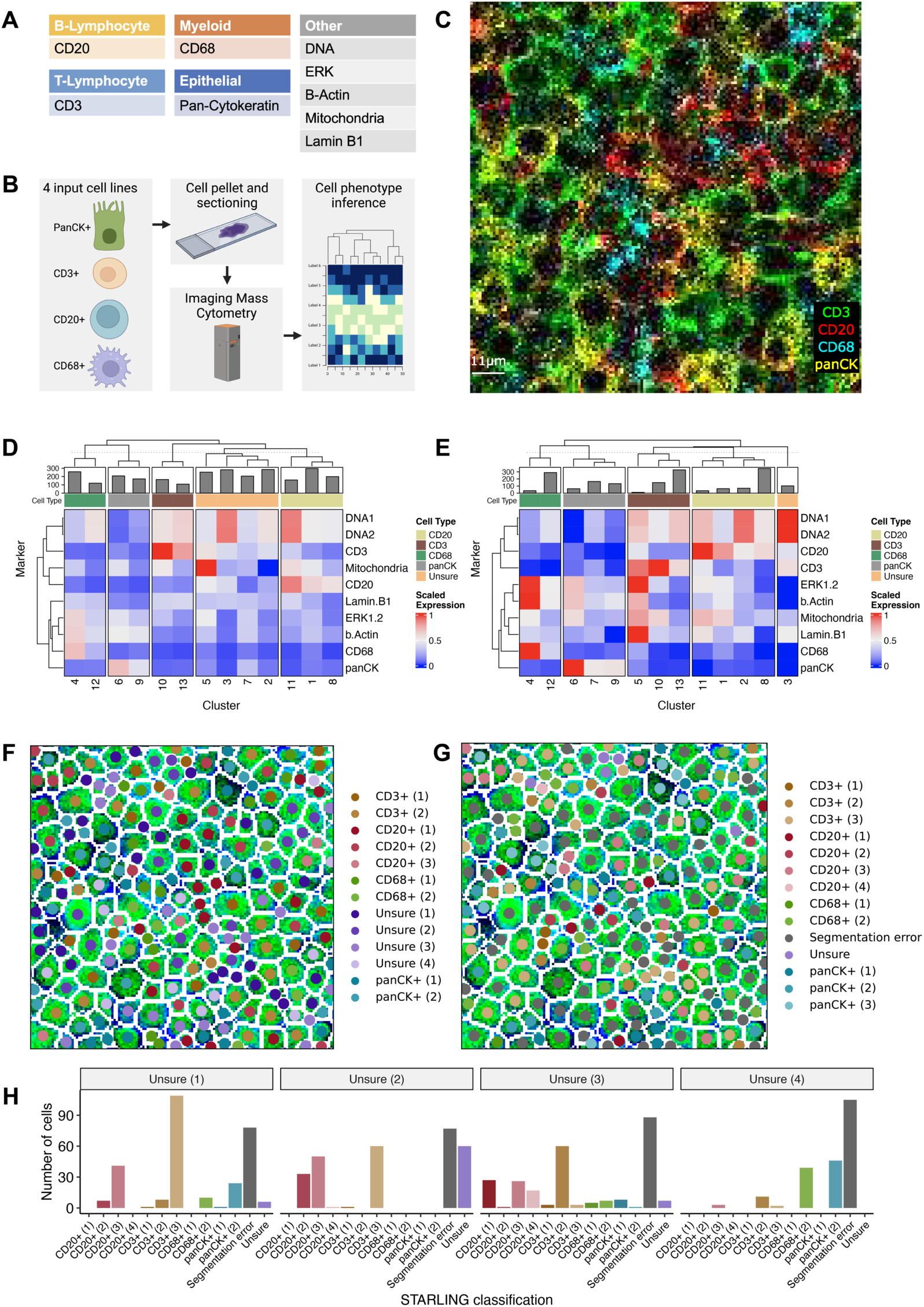
**A** Antigens targeted by the antibody panel used to profile the cell pellet data. **B** Overall workflow of IMC profiling of four cell lines to generate a controlled dataset for benchmarking. **C** Example ROI in resulting image coloured by major lineage markers. **D** Cell clusters and cell type interpretations using a standard PhenoGraph workflow. **E** Cell clusters and cell type interpretations after applying STARLING. **F** Same ROI as **(C)** coloured by PhenoGraph cell types. **G** The ROI coloured by STARLING cell type assignments. **H** The number of cells previously classified as Unsure (1-4) assigned to each cell type by STARLING.

Next, we applied STARLING using the PhenoGraph clustering for initialization. Using the same cell type assignment scheme as before, we were subsequently able to map 12/13 clusters as belonging to a given cell type (**Fig. 4E**). Quantifying the difference from the highest most expressed lineage marker to the second highest across all clusters showed an increase in discrimination for all but two of the clusters (**S. Fig. 3**), further demonstrating STARLING’s ability to recapitulate ground truth cell types in the dataset. This finding was robust to the precise threshold used to discriminate cell types (**S. Table 2**). Finally, we examined the spatial distribution of imaged cell types for both PhenoGraph (**Fig. 4G**) and STARLING (**Fig. 4H**) for the same region of interest as **Fig. 4C**. We noted that cells assigned under PhenoGraph as “unsure” largely corresponded to those identified as having segmentation errors by STARLING. To quantify this, we subsetted to all cells assigned to an unsure cluster using PhenoGraph across the entire image, and examined their assignments following STARLING (**Fig. 4H**). For 3/4 of the initially uncertain clusters the most common STARLING assignment was to a segmentation error, with segmentation errors being the second most likely assignment for the remaining cluster. This demonstrates both that segmentation errors may induce their own cell clusters in multiplexed imaging data and that STARLING may successfully guide users between biologically “real” cellular phenotypes compared to those induced primarily by segmentation errors.

### Robustly deconvoluting tissue architecture and heterogeneity in human tonsils

To demonstrate the ability of STARLING to uncover insights into tissue architecture and intra- and inter-sample cellular heterogeneity in densely packed tissues, we profiled over 240,000 cells of the human tonsil across 16 ROIs and 3 donors using an immune-focused antibody panel (**Fig. 5A, S**. **Table 3** and **Methods**). Visualization of the resulting ROIs using canonical markers clearly demonstrated known anatomical structures of the tonsil such as CD20+ germinal centers and surrounding epithelium (**Fig. 5B**). Applying STARLING to the segmented single-cell expression profiles uncovered a total of 25 clusters (**Fig. 5C** and **Methods**), which we successfully categorized into nine distinct cell types consistent with major immune and stromal lineages. This included extensive B and T cell heterogeneity consistent with the major cell populations in lymphoid tissue, including naive and activated B and T cell populations, distinct IFNγ+ and PD-1+ B cell populations, and multiple myeloid populations including macrophages, monocytes, and neutrophils. Notably, interpretation of cell clusters was improved compared to baseline methods (**S. Fig. 4**).

**Figure 5.**
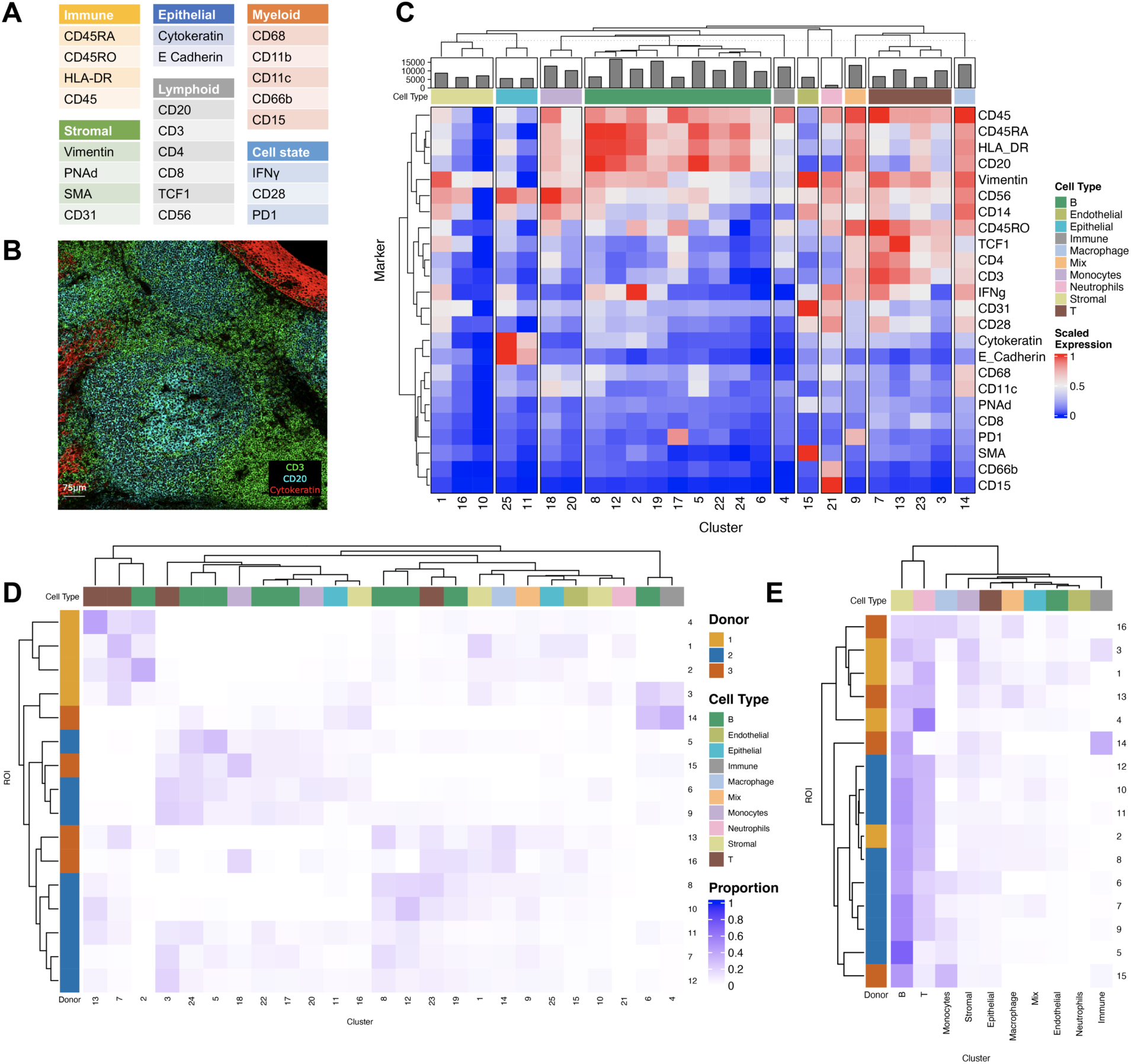
**A.** Antigens targeted in antibody panel designed for tonsil tissue. **B** Example ROI with CD3, CD20, and Cytokeratin identifies a B cell rich germinal center with peripheral T cells and tissue epithelium. **C** Mean marker expression for cluster identities discovered by running STARLING simultaneously across all 16 ROls. **D** Cluster proportions identified for each cluster in ROI highlights intra and inter-sample heterogeneity. **E** Cluster proportions for each ROI summarized to the cell type level.

Next, we quantified how cell type abundance varies between ROIs from the same donor (intra-donor heterogeneity) and between ROIs from different donors (inter-donor heterogeneity). By quantifying the proportion of each cluster in each ROI (**Fig. 5D**), we found several clusters enriched in a donor-specific manner. For example, Donor 1 is specifically enriched for clusters 2, 7, and 13, corresponding to specific populations of IFNγ+ B cells and IFNγ+ naive and activated T cells. In contrast, some clusters such as 4 and 6 (corresponding to B and other CD45+ cells) are specific to two ROIs from two distinct donors. However, when collapsing this proportion analysis to major lineage type (**Fig. 5E**), the frequencies are largely consistent across cell types showing enrichment for B and T cells as expected in the tonsil, with several exceptions. This highlights the potential of STARLING coupled with highly multiplexed imaging technologies to reverse out cell type heterogeneity in densely packed tissues.

Finally, we sought to understand how the spatial architecture of the cell types present in the tonsil varied within and between donors. We implemented a spatial enrichment analysis to identify which cell clusters are closely mixed or highly excluded spatially across the images (**Methods**). While general patterns were evident consistent with the tissue of origin such as T-B exclusion due to the spatial architecture of germinal centers, we observed extensive variability in spatial enrichment both within and between donors (**Fig. 6A**). We first checked the consistency of this spatial enrichment under two strategies for assigning cells: removing cells with a high segmentation error probability before the enrichment analysis, or using STARLING’s maximum likelihood estimate of the cell cluster assuming no segmentation error and therefore including all cells. We found the two estimates to be highly correlated (**Fig. 6B**, ρ=0.977, p<0.001), demonstrating we can robustly quantify spatial enrichment even in the presence of segmentation errors. Finally, we explored specific examples of this enrichment. For example, ROI 1 (**Fig. 6C** left) is identified as an outlier for neutrophil infiltration into the stroma compared to all other ROIs as validated by specific marker expression (**S. Fig. 5**). Conversely, ROI 11 (**Fig. 6C** right) is identified as having the most extreme compartmentalization between T and B cells.

**Figure 6.**
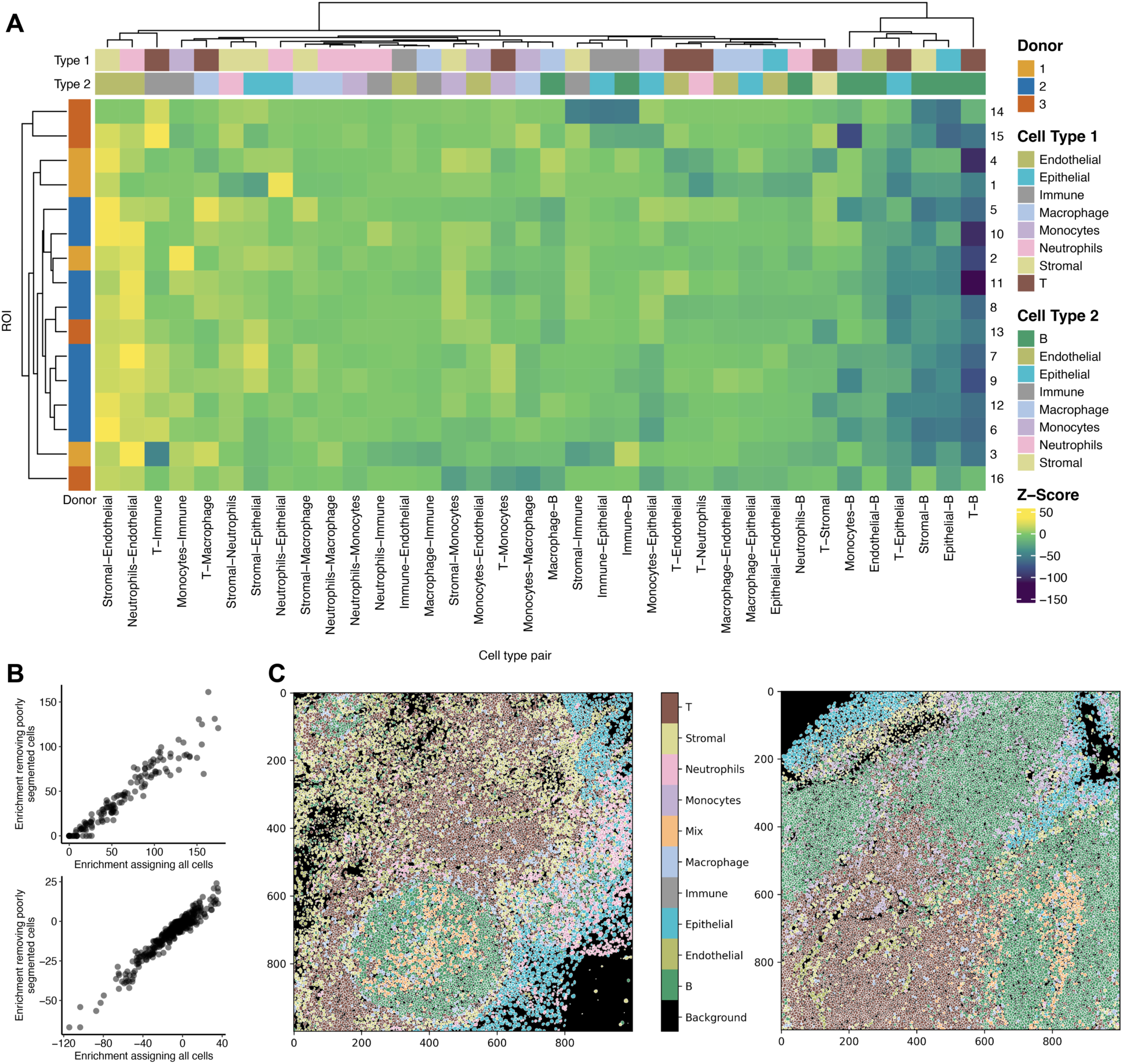
**A** Spatial enrichment of cell type pairs across all tonsil ROls. **B** Comparison of enrichment values within the same cell type (top) or between cell types (bottom) for two strategies of cluster assignment, either using all cells assigned to their most likely segmentation-error-free clusters or by removing cells with likely segmentation errors before assignment. **C** Images of ROls 1 (left) & 11 (right) coloured by inferred cell type.

## Discussion

While there is much effort in the community to improve the process of cell segmentation from multiplexed imaging^29^, fundamental physical limitations such as image resolution (>1um), signal spillover, and 3D tissue slicing effects mean a perfect segmentation is likely impossible.

Therefore, we have introduced an algorithm that explicitly acknowledges the input segmentation likely contains errors and uses knowledge of how these imperfections likely impact the quantified expression profiles to effectively denoise the data and learn the underlying cellular phenotypes. Part of the difficulty in approaching this problem was both the lack of ground truth and prior work in this area, which necessitated the development of a complete benchmarking workflow including the generation of control data as well as the creation of a plausibility score to quantify the quality of inferred cellular phenotypes.

In developing STARLING, we sought to strike a balance between practical performance and model complexity, which leads to several limitations that may serve as future directions for development. Firstly, beyond the size of each cell we do not explicitly use spatial information in the image. The number of possible reasons for a segmentation error in highly multiplexed imaging meant we opted for an approach that modeled the consequences on cellular expression rather than explicit spatial relationships giving rise to segmentation errors. While this had the advantage that we could use the consequently held-out spatial information as part of the benchmarking setup, future methods may wish to explicitly build this into the model.

Secondly, our model is mis-specified with respect to the true data generating process, assuming each segmentation error comes from the pairwise combination of cell clusters before averaging over all possible pairings to compute the final segmentation error probability (see **Methods**). We opted for this as models closer to the true data generating distribution were too flexible and required additional prior information in preliminary testing and our empirical analyses above show this mis-specified approach achieves strong performance in the absence of further *a priori* information. However, future methods may wish to explicitly consider more complex but better specified models incorporating additional prior information.

Beyond STARLING, we believe there is huge scope for segmentation-error aware methods for highly multiplexed imaging and even spatial transcriptomics. For example, though not explored here, dimensionality reduction methodologies such as UMAP^30^ are an important step in the data analytic workflow of multiplexed imaging data. Segmentation error aware dimensionality reduction could improve the ability of researchers to both interpret the data as well as integrate diverse datasets across conditions and tissues. Similarly, we previously published methodology^12^ for automated assignment of cell types in highly multiplexed imaging, where each cell is assigned to an *a priori* specified cell type based on marker over-expression. Variants of such methodology that are explicitly aware of segmentation errors would no doubt improve cellular quantification.

## Methods

### Cell pellet preparation

1 000 000 cells from each cell line (Jurkat, RAMOS, THP1, BxPC3) were pooled together and washed three times in cold PBS (4 °C). Cells were suspended in 67 μL human plasma and 116 μL bovine thrombin and allowed to coagulate for 10-15 minutes on ice. The resulting pellet was transferred into a biopsy capsule and fixed in 10% NBF for 12-24 hours, followed by 3 washes in cold PBS, 1 wash in 50% EtOH, and 1 wash in 70% EtOH for 30 minutes each. Samples were stored in fresh 70% EtOH in a fridge until paraffin embedding and microtome sectioning into 4 μm sections.

### Cell pellet and tonsil staining

Tonsil sections were formalin-fixed and paraffin embedded, and obtained from the surgical pathology unit at Mount Sinai Hospital, Toronto, Canada (REB# 20-0178-E). For IMC staining, cell pellet and tonsil sections were deparaffinized by first baking for 1 hour at 60 °C, followed by dewaxing in a series of 3 xylene solutions for 10 minutes each and then rehydration in a graded series of alcohol (ethanol:deionized water 100:0, 100:0, 96:4, 90:10, 80:20, 70:30) at 5 min each. Heat-induced epitope retrieval was conducted in a pH 9.2 Tris-EDTA buffer at 95 °C for 30 minutes. Samples were allowed to cool and blocked using a 3% BSA and 5% horse serum in 0.1% TBST (Tris buffered saline (TBS) containing 0.1% Tween-20) blocking/staining buffer for 1 hour at room temperature. Unconjugated primary antibodies were stained overnight at 4 °C, followed by conjugated secondary antibodies for 1 hour at room temperature and conjugated primaries overnight at 4 °C, with 15 minute TBS washes after each stain. All antibodies were diluted in the blocking/staining buffer. Iridium staining (^191^Ir/^193^Ir) was done at 500 nM in TBS for 5 minutes. Slides were washed for 15 minutes in TBS, dipped into ddH2O, and air dried before the IMC acquisition. Information for antibodies used to stain cell pellet and tonsil sections, including metal tag, clone, company, catalog and lot numbers, are provided in **S. Table 3 & 4.**

### Imaging mass cytometry

Images were acquired using a Hyperion or Hyperion XTi Imaging System (Standard Biotools). Chosen regions of interest on the cell pellet or tonsil sections were acquired using laser-ablation in a rastered pattern at 400 or 800 Hz, and preprocessing performed using commercial acquisition software (Standard Biotools).

### Data pre-processing for tonsil data

Tonsil IMC data were preprocessed and segmented using an in-house integrated flexible analysis pipeline (ImcPQ) available at https://github.com/JacksonGroupLTRI/ImcPQ. The analysis pipeline is implemented in Python. Briefly, data were converted to TIFF format and segmented into single cells using the pipeline to classify pixels based on a combination of antibody stains to identify membranes/cytoplasm and nuclei (**S. Table 5**). The stacks were then segmented into single-cell object masks.

### Data preprocessing for cell pellet data

Cells were segmented using the Mesmer model in DeepCell v0.12.3 using an average of the two DNA channels as the nuclear channel and an average of the four cell marker channels (b-Actin, CD20, CD3, CD68) for the cytoplasm/membrane channel. Arcsinh normalization with a cofactor of 5 was applied to all channels prior to averaging. Single-cell expression for each marker was quantified as the mean pixel value for each channel. Clustering was performed with Scanpy v1.9.5 and Phenograph v1.5.7 using Phenograph Leiden clustering on default parameters, followed by STARLING. Clusters were assigned to the cell type associated with the highest normalized expression cell type marker if the difference with the next highest cell type marker is at least 0.15.

### Data re-processing for public datasets

In this study, the Basel cohort (Jackson et al., 2020) and the Metabrics cohort (Ali et al., 2020) were selected. We utilized 358 and 548 images from the Basel and Metabrics cohorts, respectively, along with 16 unpublished Tonsil images. We employed Mesmer (Deepcell v0.12.4) for generating single-cell masks. For images from Basel and Metabrics, control channels and the markers H3K27me3, pHH3, and totHH3 were excluded. We designated DNA1 and DNA2 as nuclear channels, while the other markers were categorized as cytoplasm/membrane channels. In the case of Tonsil images, only DNA1 and DNA2 were included as nuclear channels (excluding TCF1), and the other markers were considered as cytoplasm/membrane channels. After defining the marker types, we calculated the average nuclear and cytoplasmic expression levels at each location in each image. This resulted in two distinct channels as required for Mesmer input parameters: one representing the nuclear channel and the other representing the cytoplasm channel. Lastly, we determined the single-cell expression for each protein by calculating the average expression levels within the boundaries of each mask, inclusive.

### Expression matrix

In addition to utilizing the segmentation masks provided by Basel and Metabrics cohorts, we generated our own segmentation masks using MESMER. Unlike the Basel and Metabrics cohorts, the Tonsil images lacked panel spillover corrections, requiring additional processing steps. To address the spillover issues, we obtained the spillover matrix from Jackson’s lab, which was experimentally generated. Subsequently, we applied the CATALYST R package for the necessary corrections ^15^.

Next, we utilized MESMER ^14^, a cell segmentation method for generating cell masks from multiplexed imaging data. This method is recognized for its accuracy, comparable to human performance, without the need for manual annotation, making it seamlessly integrated into our pipeline. To obtain the masks, we separately normalized (min-max) the cytoplasm/membrane and nuclear channels. Subsequently, we calculated averaged images from these channel types, as required for MESMER’s input. Following this, the two resulting images underwent another round of normalization (min-max) before being used in MESMER, generating masks that represent single cells.

To create a cell’s expression, we mapped its masked locations to the same locations in the IMC image to compute the cell expression by averaging expressions for each measured protein within each masked area (border inclusive). This process was repeated for all masks in each ROI (i.e., image), resulting in a cell expression matrix of *N* cells and *P* proteins.

### Plausibility score

We introduced plausibility scores (PS) to turn to prior knowledge of cell type markers into a quantitative measure of clustering performance. Each marker for each cluster is categorized as either non-expressed (if the average expression is in the lower 25% of all clusters) or expressed otherwise. Next, we collate a priori known sets of markers that should either never be co-expressed (e.g. T Lymphocyte marker CD3 and B Lymphocyte marker CD20) in the tissue of origin, or should always show conditional co-expression (e.g. CD3 conditional on CD45).

For each pair and each scenario, we next define “allowed” regions. For markers never expected to be co-expressed, this corresponds to the three quadrants that touch one of the axes, i.e. the regions where neither marker is expressed or only one is. For the conditional co-expression scenario where marker 1 is only expressed if marker 2 is, this region corresponds to the quadrants where (i) neither marker is expressed, marker 2 is expressed but not marker 1, or marker 1 and 2 are both expressed. The plausibility score is then simply computed as the proportion of clusters that fall into the allowed regions, averaged over all marker pairs for which information exists about co- or mutually exclusive expression.

To assess Plausibility Scores (PS), we divided cells into two distinct categories: neighboring and non-neighboring cells. A cell was classified as a neighboring cell if at least a portion of its border was adjacent to another cell. For each category, we implemented clustering algorithms— PhenoGraph, FlowSOM, and Kmeans—on their respective expression matrices, using the default settings for these clustering methods. To further evaluate the effectiveness of PS, we experimented with arcsinh transformation on the raw data and as well as using cofactors of 1 and 5. Additionally, each unique parameter setting was run 50 times, with a subsample of 10,000 cells each time. It is important to note that in cases where a category had fewer than 10,000 cells, such as with the Tonsil images, we upscaled the data with replacement.

### STARLING

For a given cell expression matrix measuring *N* cells and *P* markers, highly multiplexed imaging quantifies a vector of cell sizes 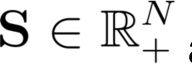 and a matrix of protein expression 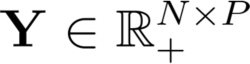. Here, we augment a mixture model to include further latent variables representing whether each cell has been segmented with errors, and if so which underlying clusters of cells contribute to the segmentation error. Let 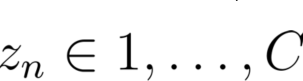 be the cluster indicator of cell *n* for one of *C* clusters. We introduce a binary variable *d_n_* that signifies whether cell *n* is a mis-segmented cell, and 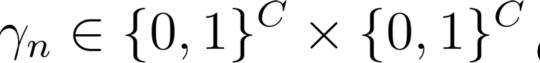 denotes the possible pairings of clusters to which segmented cell *n* belongs in the case *d_n_* = 1.

Next, we Introduce cluster centroids 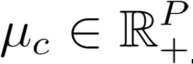. Our model assumes that conditioning on the cell containing a segmentation error and the cluster identities of each true cell being 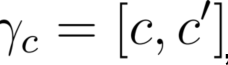, assuming a Gaussian or Student-T likelihood results in additive means and variances with averaged mean expression profiles 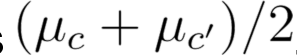. This allows us to specify the following generative probabilistic model:

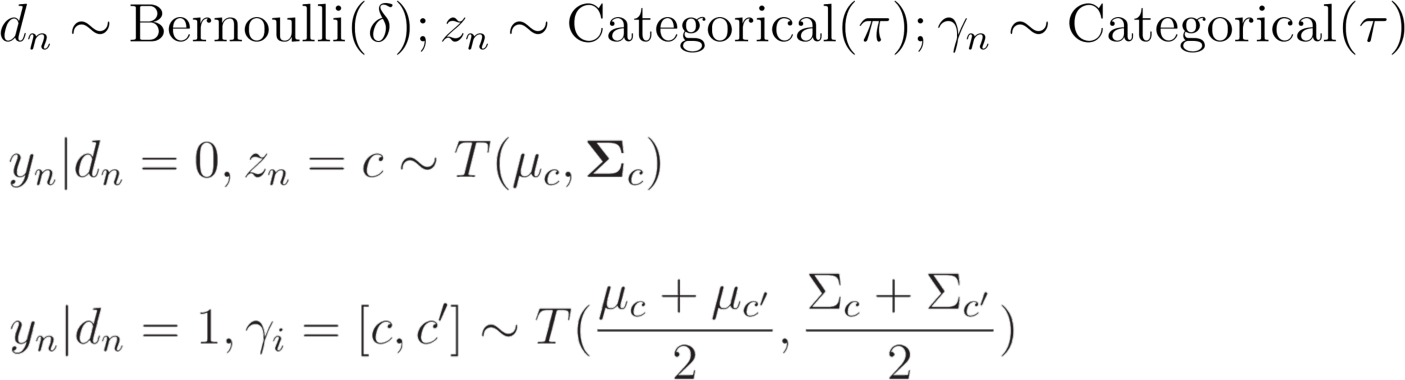

Note that the generative model here is mis-specified with respect to the true data generating distribution: the segmentation error scenario is assumed to come from equal contributions from two clusters, where in reality a segmentation error can result from any number of clusters being averaged under any proportions. We opted for this formulation as preliminary model development suggested inference under a more faithful model was too flexible and would require significant prior knowledge in order to fix what the “true” cell types were which is undesirable. Furthermore, when we compute segmentation error probabilities, we average over every possible pairing of segmentation errors weighted by their relative probability, which explains the ability to capture general segmentation errors as demonstrated above.

We further incorporate cell size information into the model. We assume in the case of a singlet, the cell size scatters around the average cell size for that cluster as 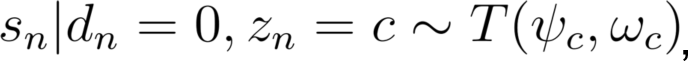, where 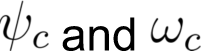 denote the cluster cell sizes and variances. However, in the case of a wrongly segmented cell, the cell size may lie in-between the maximum size of either cluster in the case of maximum z-plane overlap, to the sum of the cluster sizes in the case of no overlap. Fortunately, we can analytically integrate out this unknown cell size and defining 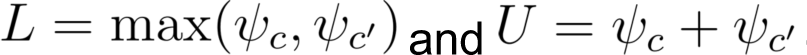 gives:

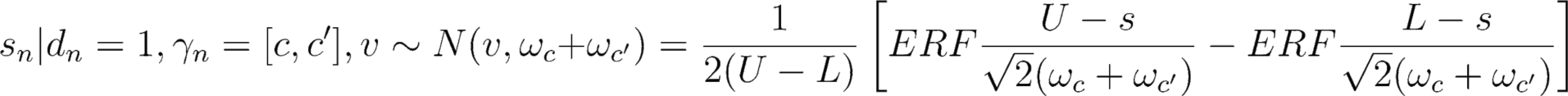

To further constrain the model, we noted that segmentation error cells may be cheaply synthesized by averaging the expression of existing cells as is common when training scRNA-seq doublet detection methods^25^. Therefore, rather than just maximize the likelihood of the model as described above, we add a discriminator to the loss that ensures the resulting probabilities 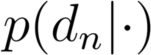 as calculated by the model can accurately identify synthetic imperfect cells vs. singlets (the overall dataset of which we denote *D̂*). Therefore, the loss function becomes:

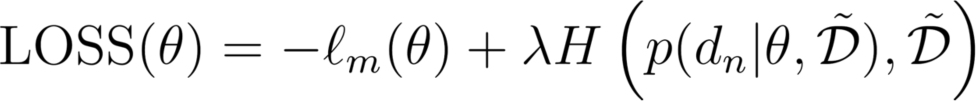

where *l_m_*(θ) is the log-likelihood function from the model, *H*(ᐧ) is a Binary Cross-Entropy (BCE) function applied to the model predicting the synthesized data, and λ is a fixed hyper-parameter that controls the balance between Log-Likelihood from the model and Binary Cross-Entropy (BCE) from the simulated data (default λ=1). Higher λ values accentuate the importance of the synthetic data (singlet detection), while lower values emphasize the input data (data likelihood) more.

Moreover, we set the segmentation error rate and γ for the synthetic data (see synthetic data) and as well as the segmentation error prior to be 40% and uniform prior for singlet (π) and segmentation error clusters (τ). STARLING is trained using Adam optimization with a learning rate of 1e-3 and minibatching (size 512) for both CPU and GPU enabled machines. For each dataset, we executed STARLING five times with random initialization, subsampling 10,000 cells each time to evaluate the model performance. After identifying the optimal set of model parameters, STARLING was then applied to each cohort without subsampling. This process was repeated five times, selecting the run with the lowest likelihood score for further analysis.

### Benchmarking with plausibility score

To evaluate cluster centroid quality, we employed compared the plausibility scores (see above) for the centroids computed by STARLING compared to three common clustering algorithms: Kmeans (as implemented in the sklearn package v0.24.2), PhenoGraph (as implemented in the scanpy package v1.7.2) and FlowSOM (v0.1.1), all using default settings. For each baseline method, we further employed a two-step approach, where a random forest algorithm (implemented using the sklearn package v0.24.2) trained to discriminate singlet cells vs. two cells pooled together, then re-predicting whether each originally segmented cell was likely a singlet, and removing cells with singlet probability < 0.5. For each analysis, we conducted an experiment involving a subsample of 10,000 cells for the three datasets, repeated five times for each parameter combination. We utilized both the provided masks and those obtained by MESMER, resulting in a total of 720 distinct model variations for each data source. For each experiment, the number of clusters (k) was initially determined using PhenoGraph, and this value was then applied consistently in Kmeans, FlowSOM and STARLING to ensure fair comparisons.

### Benchmarking for segmentation error detection

To benchmark STARLING’s ability to detect segmentation errors, we computed 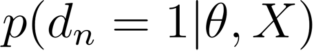 for every cell as the probability of a segmentation error. We segmented each dataset using MESMER v0.12.4 in both “whole cell” and “nucleus” mode. We denoted each segmented whole cell as containing a likely segmentation error if multiple cells segmented in nucleus only mode were found in a single cell segmented in whole cell mode. To confidently map cells, we further restricted the set of whole cells denoted as likely segmentation errors to those where at least one nuclear only segmentation took up >30% of the whole cell segmentation area.

To correlate STARLING’s ability to associate segmentation errors with adjacent cells, we fit the model to estimate cluster assignments for every cell. For every cell n, we found the segmentation error free cluster identity (i.e. the max of 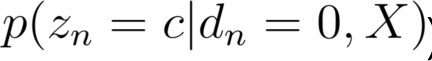) for both that cell, and all of its immediate neighbours denoted 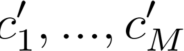 for *M* touching cells. Next, we extract the *C*x*C* matrix Γ for each *n* denoted *G* (where we drop the *n* subscript for future clarity). Next, we let *v* be the c^th^ row of *G*, i.e. 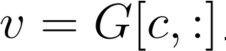 — the set of segmentation error probabilities for cell *n* for the mixing between the assigned cell type *c* and all others. Then, let *v_α_* be the elements of *v* belonging to the identities of the touching cells/neighbours, i.e. 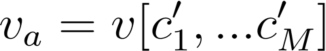 and let *v_b_* be the other elements of *v*, i.e. 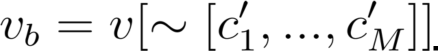. We repeated this for every cell, and the difference between *v_α_* and *v_b_* (i.e. mixing probabilities for touching vs non-touching cells).

### Tonsil analysis

STARLING was fitted using mini-batching (batch size 512) and C=25 clusters on all cells from the tonsil data ROIs for 5 different initializations, with the final model chosen by the fit that minimized the log-likelihood score. For enrichment analysis we employed the Python Squidpy package v1.2.0 to generate z-scores for a cell-type-by-cell-type square enrichment matrix. The diagonal and off-diagonal z-scores represent the z-scores of cell types within the same and different pairs of cell types, respectively. Each Region of Interest (ROI) was analyzed independently, and the results for the two cases were presented separately. Segmentation-error-free cell phenotypes analyzed with STARLING may be interpreted in two ways. Firstly, cells with high probability of segmentation error (i.e. 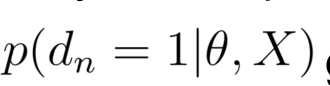 greater than 50%) may be removed, then the most likely cluster assigned (*Method 1*). Alternatively, we can ask STARLING for the “best guess” assuming a cell has no segmentation error (i.e. 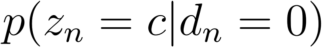), which we refer to as *Method* 2.. We assigned cells using both of these methods, with the subsequent enrichment analysis comparing them shown in **Fig. 6B**.

## Data and code availability

The STARLING Python package is available at https://github.com/camlab-bioml/starling. Code to reproduce the analyses in this paper is available at https://github.com/camlab-bioml/starling_plots.

## Acknowledgements

This work is funded by NSERC Discovery Grants RGPIN-2020-04083 to KRC and RGPIN-2021-03404 to HWJ, and a CIHR project grant PJT-175270 to KRC. Both KRC and HWJ acknowledge support from the Canada Research Chairs program and the Canadian Foundation for Innovation. This work was further supported by a University of Toronto Data Sciences Institute Research Software Development Support.

## Ethics approval

The data generation and analysis in this manuscript is covered under REB #20-0178-E at Mount Sinai Hospital, Toronto ON, Canada.

## Contributions

Conceptualization: YL, AS, HWJ, KRC. Data analysis: YL, DC, SA, KRC. Software development: YL, CK, KRC. Data generation: EC, AD, MM. Manuscript writing and review: YL, AS, HWJ, KRC.

