## Supplementary figures for "Segmentation aware probabilistic phenotyping of single-cell spatial protein expression data"

Supplementary Figure 1. Scatterplot of CD3 and CD20 across three IMC datasets.

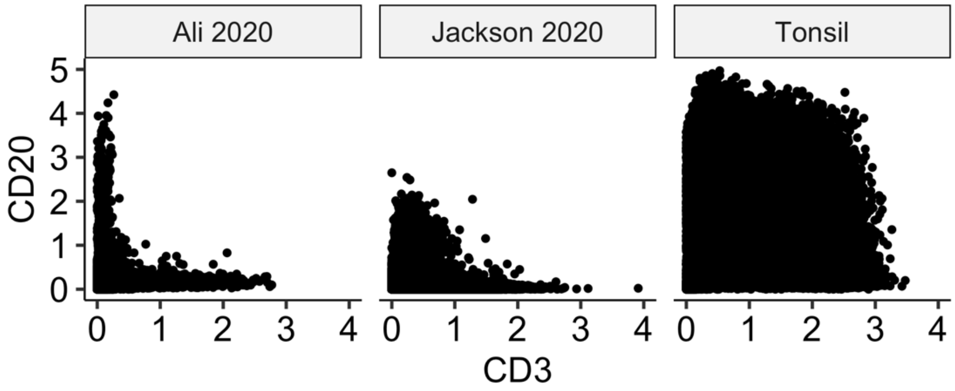


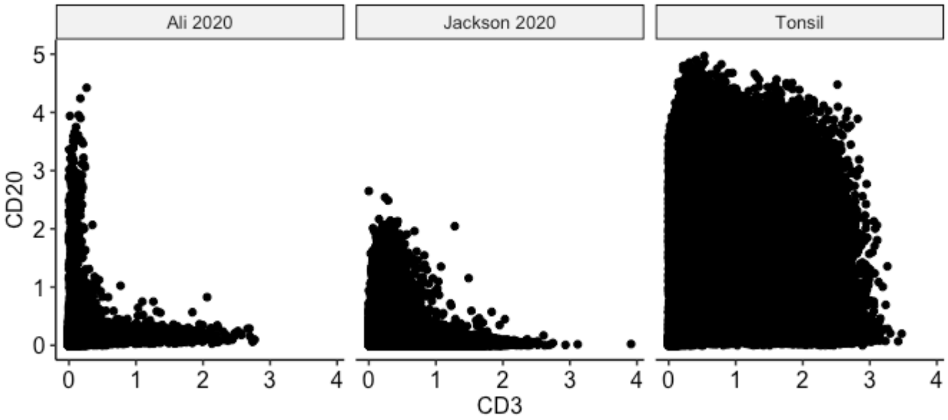


Supplementary Figure 1. Scatterplot of CD3 and CD20 across three IMC datasets.

Supplementary Figure 2. STARLING predictions analysis of segmentation error vs free and their convex area to cell area ratios.

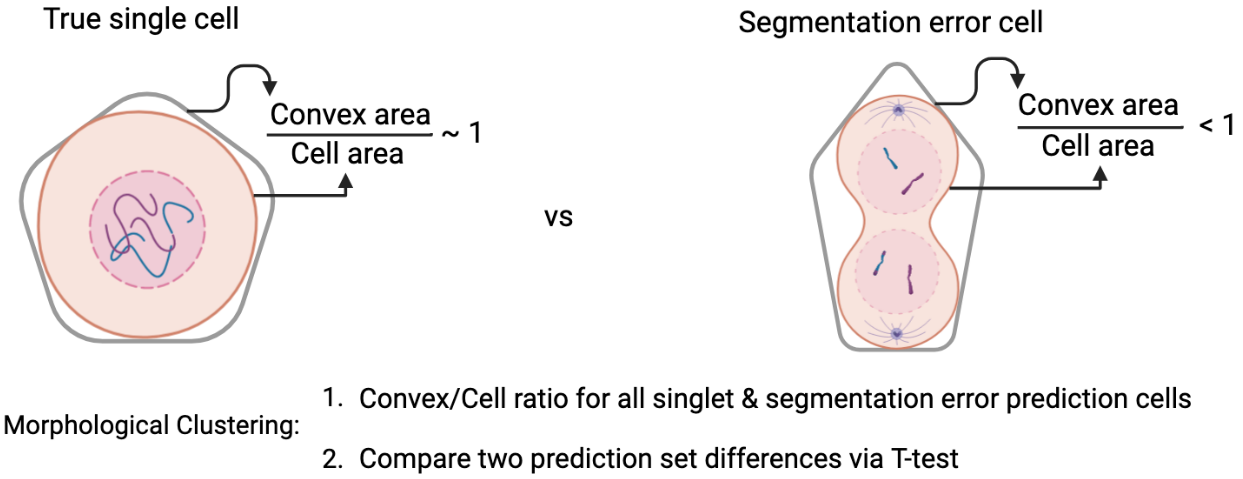


Supplementary Figure 3. The difference in expression between the highest and second highest expressed cell lineage marker of each cluster in the cell pallet data, with “Change” quantifying the difference in this quantity between the initial (PhenoGraph) and STARLING.

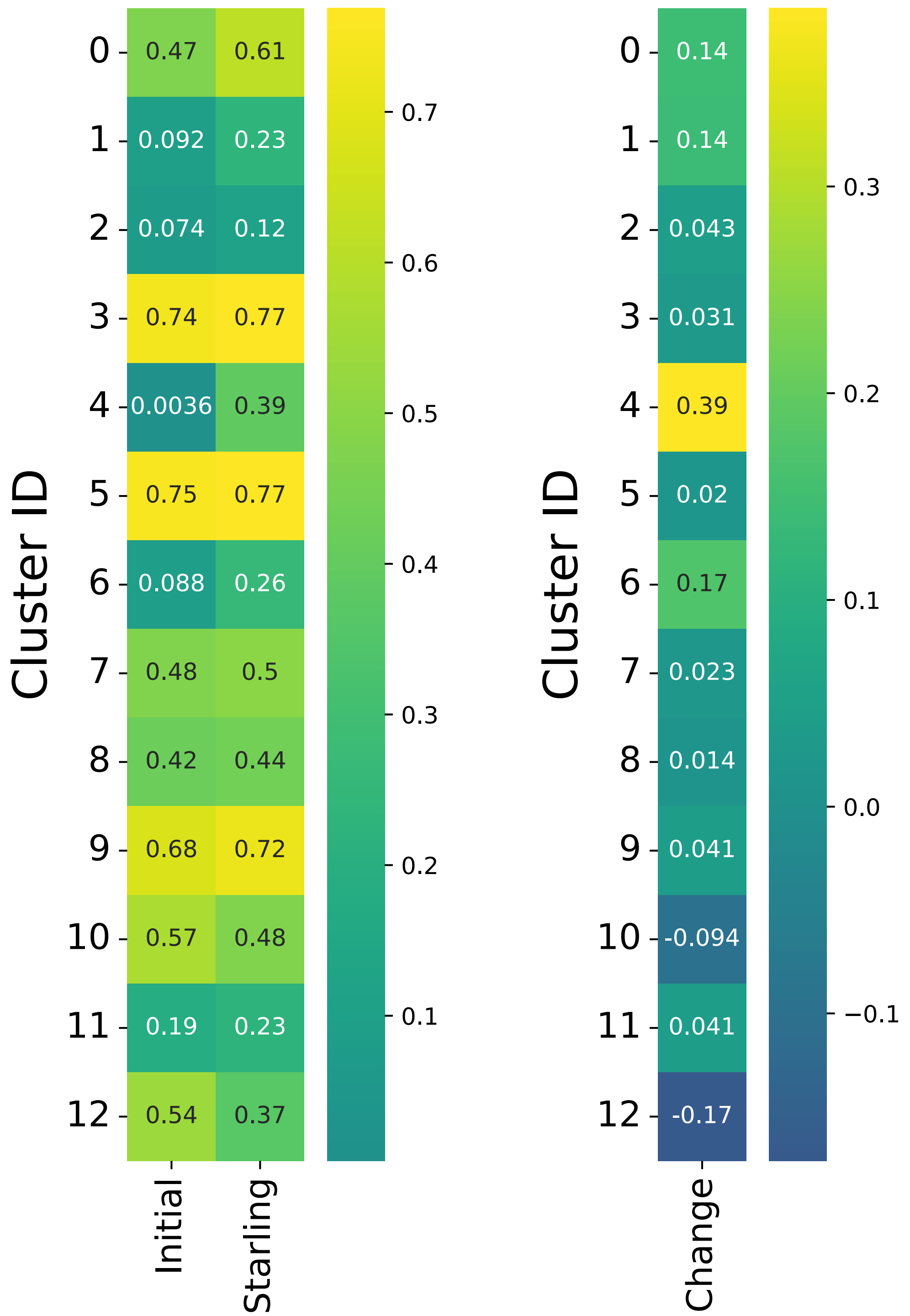


Supplementary Figure 4. Baseline method heatmaps (PhenoGraph, FlowSOM & Kmeans). CD3 and CD20 exhibit lower expression levels in B and T cell clusters, respectively, when compared to PhenoGraph on average. Additionally, PNAd expression in the Neutrophil cluster is higher in PhenoGraph.

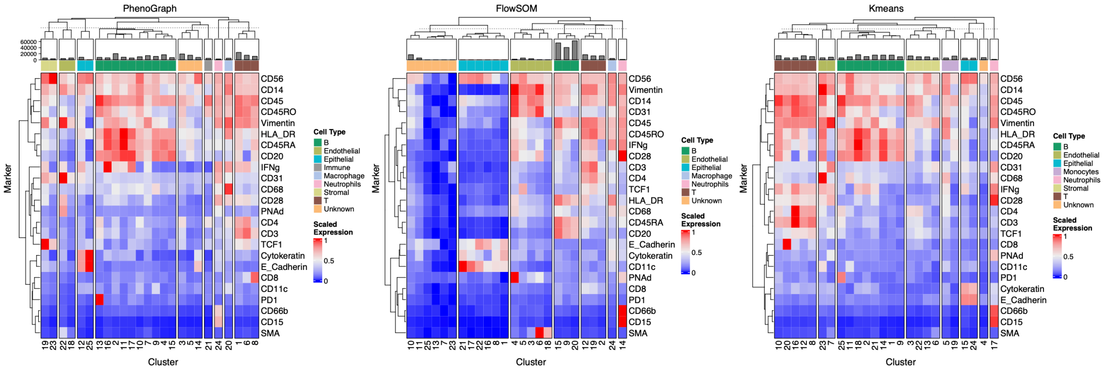


Supplementary Figure 5. CD15, CD66b and ECadherin expressions

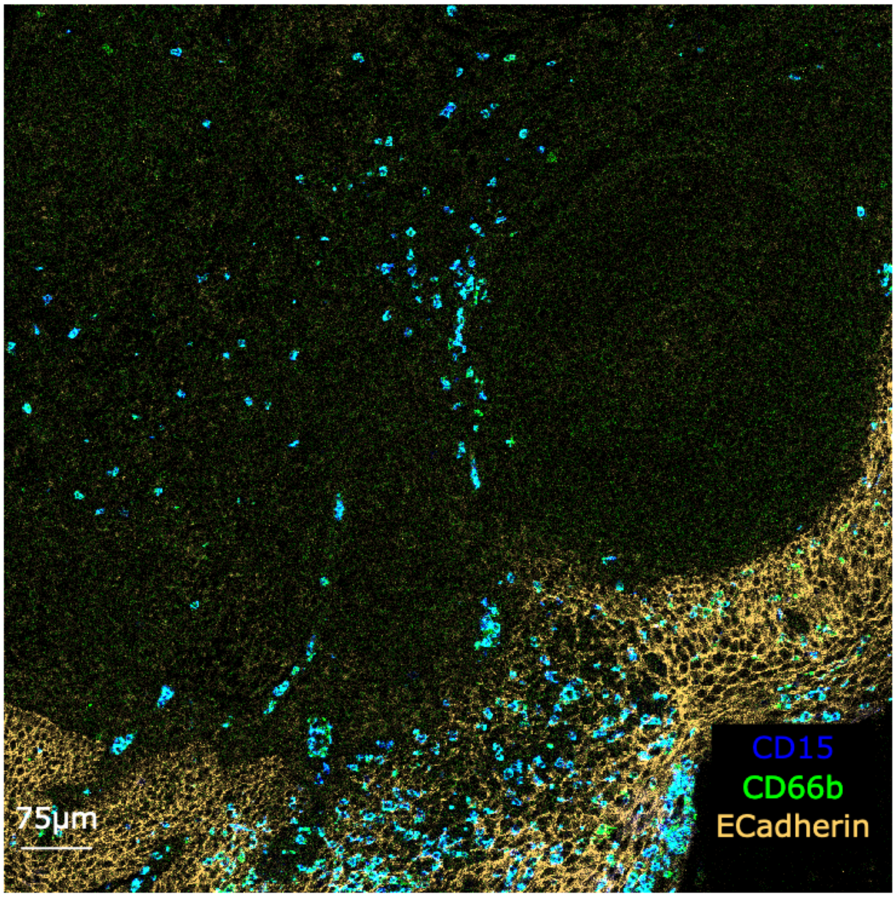
