## Supplementary tables for "Segmentation aware probabilistic phenotyping of single-cell spatial protein expression data"

Supplementary Table 1. Marker pairs used for plausibility score calculations

| Datasets | Pair Type | Marker 1 | Marker 2 |
|---|---|---|---|
| Jackson 2020/Ali 2020 | Conditional | CD45 | CD3 |
| Jackson 2020/Ali 2020 | Conditional | CD45 | CD20 |
| Jackson 2020/Ali 2020 | Conditional | CD45 | CD68 |
| Jackson 2020/Ali 2020 | Conditional | panCK | CK5 |
| Jackson 2020/Ali 2020 | Conditional | panCK | CK7 |
| Jackson 2020/Ali 2020 | Conditional | panCK | CK8&18 |
| Jackson 2020/Ali 2020 | Conditional | panCK | CK19 |
| Jackson 2020/Ali 2020 | Mutually exclusive | CD3 | CD20 |
| Jackson 2020/Ali 2020 | Mutually exclusive | CD3 | CD31 |
| Jackson 2020/Ali 2020 | Mutually exclusive | CD3 | CD68 |
| Jackson 2020/Ali 2020 | Mutually exclusive | CD3 | ECadherin |
| Jackson 2020/Ali 2020 | Mutually exclusive | CD3 | panCK |
| Jackson 2020/Ali 2020 | Mutually exclusive | CD20 | CD31 |
| Jackson 2020/Ali 2020 | Mutually exclusive | CD20 | CD68 |
| Jackson 2020/Ali 2020 | Mutually exclusive | CD20 | ECadherin |
| Jackson 2020/Ali 2020 | Mutually exclusive | CD20 | panCK |
| Jackson 2020/Ali 2020 | Mutually exclusive | CD31 | CD68 |
| Jackson 2020/Ali 2020 | Mutually exclusive | CD31 | ECadherin |
| Jackson 2020/Ali 2020 | Mutually exclusive | CD45 | panCK |
| Jackson 2020/Ali 2020 | Mutually exclusive | CD68 | ECadherin |
| Jackson 2020/Ali 2020 | Mutually exclusive | panCK | Vimentin |
| Tonsil | Conditional | CD3 | CD4 |
| Tonsil | Conditional | CD3 | CD8 |
| Tonsil | Conditional | CD45 | CD3 |
| Tonsil | Conditional | CD45 | CD4 |
| Tonsil | Conditional | CD45 | CD8 |
| Tonsil | Conditional | CD45 | CD20 |
| Tonsil | Conditional | CD45 | CD68 |
| Tonsil | Mutually exclusive | CD3 | CD20 |
| Tonsil | Mutually exclusive | CD3 | CD31 |
| Tonsil | Mutually exclusive | CD3 | CD68 |
| Tonsil | Mutually exclusive | CD3 | ECadherin |
| Tonsil | Mutually exclusive | CD4 | CD8 |
| Tonsil | Mutually exclusive | CD4 | CD20 |
| Tonsil | Mutually exclusive | CD4 | CD31 |
| Tonsil | Mutually exclusive | CD4 | ECadherin |
| Tonsil | Mutually exclusive | CD8 | CD20 |
| Tonsil | Mutually exclusive | CD8 | CD31 |
| Tonsil | Mutually exclusive | CD8 | CD68 |
| Tonsil | Mutually exclusive | CD8 | ECadherin |
| Tonsil | Mutually exclusive | CD20 | CD31 |
| Tonsil | Mutually exclusive | CD20 | CD68 |

| | | | |
|---|---|---|---|
| Tonsil | Mutually exclusive | CD20 | ECadherin |
| Tonsil | Mutually exclusive | CD31 | CD68 |
| Tonsil | Mutually exclusive | CD31 | ECadherin |
| Tonsil | Mutually exclusive | CD68 | ECadherin |

Supplementary Table 2. Threshold used for discriminate cell types

| **Identifiability Threshold** | **Number of unidentifiable initial clusters** | **Number of unidentifiable STARLING clusters** |
|---|---|---|
| 0.1 | 4 | 0 |
| 0.15 | 4 | 1 |
| 0.2 | 5 | 1 |

Supplementary Table 3. Antibody information for tonsil section stains.

| **Marker** | **Metal** | **Clone** | **Company** | **Catalog #** | **Lot #** |
|---|---|---|---|---|---|
| SMA | Y89 | 1A4 | Thermo | 14-9760-82 | 2288516 |
| E-Cadherin | In113 | 36/E-Cadherin | BD | 610182 | 1040966 |
| Pan-Cytokeratin | In115 | AE1 | Sigma | MAB1612 | 3460341 |
| Pan-Cytokeratin | In115 | AE3 | Sigma | MAB1611 | 3382323 |
| HLA-DR | Pr141 | TAL 1B5 | Abcam | ab176408 | GR3384096-1 |
| Vimentin | Nd142 | EPR3776 | Abcam | ab193555 | GR3396611-7 |
| CD28 | Nd144 | EPR22076 | Abcam | ab243557 | GR3358395-2 |
| CD15 | Nd145 | HI98 | BioLegend | 301902 | B265372 |
| CD45RA | Nd146 | HI100 | Thermo | 14-0458-82 | 2181694 |
| CD66b | Sm147 | G10F5 | BioLegend | 305102 | B349411 |
| CD20 | Sm149 | L26 | Thermo | 14-0202-82 | 2172592 |
| CD68 | Nd150 | KP1 | BioLegend | 916104 | B283618 |
| CD4 | Eu151 | EPR6855 | Abcam | ab181724 | 1023375-4 |
| CD8 | Sm152 | C8/144B | BioLegend | 372902 | B306201 |
| CD11c | Sm154 | EP1347Y | Abcam | ab216655 | GR3357092-4 |
| CD45RO | Dy162 | 55618BF | Cell Signaling Technology | UCHL1 | 2 |
| CD3 | Unconjugated | CD3-12 | Thermo | MA5-16622 | WG3333396 |
| Rat IgG | Dy163 | Polyclonal | Thermo | B305020 | 61-172-060320 |
| IFNg | Tm169 | IFNG/466 | Abcam | ab218890 | GR3424660-1 |

| | | | | | |
|---|---|---|---|---|---|
| TCF1 | Er170 | C63D9 | Cell Signaling Technology | 85942SF | 2 |
| CD14 | Yb172 | SP192 | Abcam | ab230903 | GR3273650-5 |
| CD56 | Yb173 | EPR2566 | Abcam | ab214436 | GR3317476-5 |
| PD1 | Yb176 | D4W2J | Cell Signaling Technology | 63815SF | 7 |
| CD45 | Pt194 | D9M8I | Cell Signaling Technology | 47937SF | 11 |
| PNAd | Pt195 | MECA-79 | BioLegend | 120802 | B305020 |
| CD31 | Pt196 | RM1006 | Abcam | ab282746 | GR3389585-7 |

Supplementary Table 4. Antibody information for cell pellet stains

| **Marker** | **Metal** | **Clone** | **Company** | **Catalog #** | **Lot #** |
|---|---|---|---|---|---|
| Pan-Cytokeratin | In115 | C11 | Thermo Fisher | MA1-12594 | YH4021972 |
| Pan-Cytokeratin | In115 | AE1 | Sigma Aldrich | MAB1612 | 4002485 |
| Lamin B1 | Nd143 | polyclonal_ab16048 | Abcam | ab16048 | 1042700-1 |
| b-Actin | Nd148 | SP124 | Abcam | ab242387 | GR3454668-1 |
| CD20 | Sm149 | L26 | Thermo Fisher | 14-0202-82 | 2526316 |
| CD68 | Nd150 | KP1 | BioLegend | 916104 | B373258 |
| CD68 | Unconjugated | D4B9C | Cell Signaling Technology | 26042SF | 2 |
| Rat IpG | Sm152 | Polyclonal_A18873_Thermo | Thermo Fisher | A18873 | 61-172-060320 |
| Mitochondria | Gd155 | 113-1 | Abcam | ab92824 | 1012031-1 |
| ERK (1/2) | Dy161 | 137F5 | Cell Signaling Technology | 68303SF | 1 |
| CD3 | Unconjugated | CD3-12 | Thermo Fisher | MA5-16622 | YE3922415 |
| Rabbit IpG | Yb171 | Polyclonal_A27033_Thermo | Thermo Fisher | A27033 | RL246119A |

Supplementary Table 5. Makers used for MESMER's Cytoplasm/Membrane computation

| Marker | Basel | Metabrics | Tonsil |
|---|---|---|---|
| B-Catenin | x | x | |
| CAIX | x | x | |
| CD3 | x | x | x |
| CD4 | | | x |
| CD8 | | | x |
| CD11c | | | x |
| CD14 | | | x |
| CD15 | | | x |
| CD20 | x | x | x |
| CD28 | | | x |
| CD31 | x | x | x |
| CD44 | x | x | |
| CD45 | x | x | x |
| CD45RA | | | x |
| CD45RO | | | x |
| CD56 | | | x |
| CD66b | | | x |
| CD68 | x | x | x |
| CK5 | x | x | |
| CK7 | x | x | |
| CK14 | x | x | |
| CK8/18 | x | x | |
| CK19 | x | x | |
| cMyc | x | x | |
| cPARP/cCasp3 | x | x | |
| Cytokeratin | | | x |
| ECadherin | x | x | x |
| EGFR | x | x | |
| EpCAM | x | x | |
| ER | | x | |
| ERK | x | x | |
| Fibronectin | x | x | |
| GATA3 | x | x | |
| HER2 | x | x | |
| HLA-DR | | | x |
| IFNg | | | x |
| KI67 | x | x | |
| mTOR | x | x | |
| p53 | x | x | |

| | | | |
|---|---|---|---|
| panCK | x | x | |
| PD1 | | | x |
| PNAd | | | x |
| PR | x | x | |
| RabbitIgGHL | x | | |
| S6 | x | x | |
| Slug | x | x | |
| SMA | x | x | x |
| SOX9 | x | x | |
| Twist | x | x | |
| Vimentin | x | x | x |

Supplementary Table 6. Summary of each cohort

| | Number of ROIs | Number of Cells (Provided) | Number of Cells (MESMER) |
|---|---|---|---|
| Basel (Jackson et al., 2020) | 358 | 802591 | 805675 |
| Metabrics (Ali et al., 2020) | 548 | 536883 | 589992 |
| Tonsil (Unpublished) | 16 | - | 243738 |
